# COMPASS Family Histone Methyltransferase ASH2L Mediates Corticogenesis via Transcriptional Regulation of Wnt Signalling

**DOI:** 10.1101/413674

**Authors:** Liang Li, Xiangbin Ruan, Chang Wen, Pan Chen, Wei Liu, Liyuan Zhu, Pan Xiang, Xiaoling Zhang, Qunfang Wei, Lin Hou, Bin Yin, Jiangang Yuan, Boqin Qiang, Pengcheng Shu, Xiaozhong Peng

**Affiliations:** The State Key Laboratory of Medical Molecular Biology, Neuroscience Center, Medical Primates Research Center and Department of Biochemistry and Molecular Biology, Institute of Basic Medical Sciences, Chinese Academy of Medical Sciences, School of Basic Medicine Peking Union Medical College, Beijing 100005, China; Institute of Medical Biology Chinese Academy of Medical Sciences, Chinese Academy of Medical Science and Peking Union Medical College, Kunming 650118, China; Department of Anatomy and Histology, Institute of Basic Medical Sciences, Chinese Academy of Medical Sciences, School of Basic Medicine Peking Union Medical College, Beijing 100005, China

**Keywords:** Ash2L, H3K4me3, neural progenitor cell, Wnt signalling, Corticogenesis

## Abstract

Cell fate specification in neural progenitor cells (NPCs) is orchestrated via extrinsic and intrinsic molecular programs, and histone methylation in these decisions has been ascribed to a crucial function regulating gene expression. Here, we show that the COMPASS family histone methyltransferase co-factor ASH2L is required in NPCs proliferation and upper layer cortical projection neurons production and position. Deletion of Ash2l impairs trimethylation of H3K4 and transcriptional machinery specifically for subsets of Wnt-β-catenin signalling, disrupting their transcription and consequently inhibiting the proliferation ability of NPCs in late stages of neurogenesis. Consistently, Ash2l conditional mutants exhibit thinning neocortex with reduced upper layer neurons and altered neuronal position. Moreover, overexpressing β-catenin after Ash2l elimination or knockdown can rescue the proliferation deficiency of NPCs both *in vivo* and *in vitro*. These results demonstrate an essential and highly specific role for Ash2l in controlling NPCs proliferation and late-born neurons lamination in corticogenesis via transcriptionally regulating Wnt-β-catenin signalling, and provide clues to how the COMPASS family epigenetic factors coordinate cell fate determination during cortex development.

## Introduction

The neocortex comprises an enormous variety of neuronal subtypes and glial cells that are responsible for higher order cognitive functioning and sensory processing(Jabaudon, 2017, Rakic, 2009). Assembly of a functional neocortex relies on the timely establishment of cell type-specific transcriptional programs to induce NPCs to produce the correct number and diversity of neurons for intricate circuit assembly. Radial glia cells (RGCs), the prototypical neurogenic NPCs, reside in the ventricular zone (VZ) of the cortical epithelium, initially expanding via symmetric division and then undergoing asymmetrical divisions to give rise to neurons and glia(Kriegstein & Alvarez-Buylla, 2009, Rakic, 2009). At the onset of neurogenesis, NPCs in the dorsal telencephalon undergo neurogenic divisions to produce glutamatergic projection neurons (PNs) that migrate radially to the cortical surface to generate the six layers of the neocortex. PNs are generated in a tightly regulated, inside-out order, with later generated neurons bypassing early-born cells to settle at progressively more superficial layers(Breunig, Haydar et al., 2011, Kwan, Sestan et al., 2012). The PNs distributed within each layer have different shapes, patterns of connectivity, physiological characteristics, and specific gene expression patterns(Harris & Shepherd, 2015, Jabaudon, 2017). Indeed, inappropriate gene regulatory circuitry and cell fate commitment lead to a large number of neurological disorders(Kwan et al., 2012). Therefore, it is crucial to understand the regulatory mechanisms to define distinct cell types during neocortical development.

It is widely accepted that the lineage fate determination of neocortical NPCs is a highly regulated program involving both intrinsic and extrinsic factors. Multiple signalling pathways, such as Wnt signalling, are required during distinct developmental windows to direct patterning, specification, and proliferation of progenitors(Backman, Machon et al., 2005, Draganova, Zemke et al., 2015, Munji, Choe et al., 2011). However, the process of cell fate establishment is accompanied by cell epigenetic state changes and class-specific gene expressions. Transcription factors and epigenetic modifications are increasingly being implicated in governing the intricate temporal and spatial regulation of the fate choice of NPCs(Hirabayashi & Gotoh, 2010, Imayoshi & Kageyama, 2014, Mitrousis, Tropepe et al., 2015). Post-translational modification of core histones can impact gene expression by altering chromatin structure. Tri-methylation of lysine-9 and lysine-27 in histone H3 (H3K9me3, H3K27me3) are typically marks of transcriptional silencing, and tri-methylation of lysine-4 in histone H3 (H3K4me3) is a generally connected with open chromatin and gene expression. H3K27 methylation is catalysed predominantly by Polycomb group proteins (PcG), while H3K4 methylation is mainly mediated by Trithorax group proteins (TrxG) or its mammalian homologues mixed-lineage leukaemia (MLL1-4 or named lysine-specific methyltransferases subclass 2 (KMT) 2A-D) and SET1A/B (KMT2F and G) enzymes of the COMPASS (complex of proteins associated with Set1) family(Piunti & Shilatifard, 2016, Schuettengruber, Bourbon et al., 2017). Interestingly, promoters of a subset of key developmental regulatory genes are associated with both active H3K4me3 and repressive H3K27me3 marks, also referred to as “bivalent” modification signatures, which retain the stem and progenitor cells in a poised transcriptional state and enable rapid activation or repression once a specific fate is committed(Bernstein, Mikkelsen et al., 2006). There is little known about the effects on lineage fate decision of NPCs when these bivalent modifications disrupted during neurogenesis. Recent epigenome profiling studies in NPCs have uncovered that dynamic H3K27me3 changes during neurogenesis coincide with cell fate transitions(Albert, Kalebic et al., 2017). It remains unclear how dynamic H3K4me3 changes is in the context of the cell fate determination during neocortex development. In addition to H3K27me3, multiple PcG members are important for the balance between self-renewal and differentiation during cortex development(Bruggeman, Valk-Lingbeek et al., 2005, Pereira, Sansom et al., 2010), but the function of the histone-modifying COMPASS family member in neurogenesis remains unknown.

ASH2L (Absent, Small or Homeotic 2-like) together with WDR5 (WD repeat domain 5), RbBP5 (Retinoblastoma binding protein 5), and DPY30 (Dumpy-30) (or WRAD for short) collectively form the ‘core complex’ that serves as a modulatory platform capable of efficient enzymatic activity(Dou, Milne et al., 2006, Mersman, Du et al., 2012, Shilatifard, 2012, Steward, Lee et al., 2006). Furthermore, the WRAD complex interacts with different types of KMT2 enzymes and other additional subunits that are not shared among all to form different functional complexes(Piunti & Shilatifard, 2016, Shilatifard, 2012) and these different COMPASS complexes show unique functionalities; for example, SETD1A/B complexes likely serve for global H3K4me3 deposition, whereas MLL2 complexes appear to be responsible for H3K4 trimethylation at bivalent promoters(Ardehali, Mei et al., 2011, Denissov, Hofemeister et al., 2014, Wu, Wang et al., 2008). Therefore, it is a good strategy to conditionally delete the core subunits of the WRAD complex to reveal the requirement of the COMPASS proteins and H3K4me3 for mouse development. In this study, we investigated that effect in neocortical development by conditional knockout of Ash2l. Ash2l has shown to be essential for proper H3K4 trimethylation *in vitro* (Dou et al., 2006, Steward et al., 2006). *In vitro*, Ash2l was previously shown to control embryonic stem cell maintenance and differentiation (Stoller, Huang et al., 2010, Wan, Liang et al., 2013). In the nervous system, Ash2l has also been described as a regulator of dendrite morphology in brain development(Jung, Hsieh et al., 2016). In particular, mutations in Ash2l may associate with intellectual disability in humans(Karaca, Harel et al., 2015). However, the functions of Ash2l involved in the establishment of specific transcriptional programs defining NPC lineage fates during cortical development remain largely unknown.

Here, we found that conditional deletion of Ash2l by D6-Cre resulted in hypoplasia of the developing cortex and thinning of the cortical plate, with dysgenesis of NPCs proliferation in the context of rapid cellular transitions during corticogenesis. Loss of Ash2l promotes NPCs exhaustion and cell cycle exit at late neurogenesis. Inactivation of Ash2l results in downregulated expression of neurogenesis and Wnt-β-catenin signalling-related genes via repressing tri-methylation of H3K4. And over expressing β-catenin rescued above perturbations. Together, these observations provide insights into temporal regulatory mechanisms mediated by COMPASS family epigenetic factor Ash2l as a transcriptional activator that maintains the NPCs pool and proliferative competence during late stages of neocortical development.

## Results

### Dysregulation of the Laminar Structure and Neuron Specification in the Ash2l cKO Postnatal Cortex

To understand the role of Ash2l in cortical development, we first analysed the spatial and temporal expression pattern of *ash2l* mRNA by *in situ* hybridization with an *ash2l* anti-sense probe. We chose embryonic day (E) 12.5, E14.5 and E16.5 as the three neurogenic stages of the developing mouse brain to examine. At all three stages, Ash2l is widely expressed in the cerebral cortex, with high expression in the progenitor zone (Appendix Fig S1A). Furthermore, the open access, single-cell RNA-sequencing (RNA-seq) dataset of RGCs and their progeny(Telley, Govindan et al., 2016) revealed specific Ash2l expression in intermediate progenitors (IPs) and late-born neurons, suggesting that Ash2l may play specific roles in NPC fate determination during neurogenesis (Appendix Fig S1B).

Then, we generated Ash2l conditional knockout mice using the Cre/loxP system and chose the D6-cre knock-in mouse line in which the D6 promoter/enhancer starts to be activation in the dorsal cortex at E10.5(van den Bout, Machon et al., 2002) to mediate Ash2l deletion in the early neocortical developmental stage (Appendix Fig S1C). At E14.5, Ash2l expression was nearly depleted in the cerebral cortex (Appendix Fig S1D). Ash2l^fl/fl^;Cre^+^ (termed Ash2l cKO) mice were born in Mendelian ratios but experienced significant growth retardation and infertility, while the littermate Ash2l^fl/+^; Cre^+^ mice and Ash2l^+/+^ Cre^+^ (termed Control) did not show detectable phenotypes. We quantified the survival of Ash2l cKO mice that were genotyped at 1 week of age and tracked over 12 months. Almost half of the Ash2l cKO mice died around 1month age (Appendix Fig S1E). These mice exhibited hydrocephalus (Appendix Fig S1F) and spontaneous epileptiform activity (Video S1). Mutant mice also showed thinner lateral cortex and enlargement of the lateral ventricle (Appendix Fig S1G-1H). Thus, Ash2l function in the nervous system is essential for life in mammals.

To further examine of the neuronal organization of the Ash2l cKO neocortex, we performed immunostaining for upper layer PN markers Brn2 and Cux1 and deep layer PN markers Ctip2 and Tle4. We focused our analysis on the dorsal cortex (boxed area in Fig 1A and 1B) at P3 and found that a marked difference in cellular composition, both the location and number, of cortical PNs were abnormal in the Ash2l cKO neocortex (Fig 1C). We observed significant reduction of Brn2-and Cux1-positive cell numbers (Fig 1E). In addition, Brn2-and Cux1-positive cells of Ash2l cKO mice were visibly misplaced in deep cortical layers and in the underlying white matter (Fig 1C). Displaced Cux1 cells found in the deep cortical layers maintained their molecular identity as superficial layer PNs, as all of them expressed the superficial layer-specific marker Cux1 but were not labelled with Ctip2 or Tle4 (Fig 1D). These observations suggested a deficiency in the production and position of PNs that typically occupy the upper cortical layers. For the deep layer cortex, the layer V marker Ctip2 was significantly reduced; it appeared to be replaced by Tle4^+^ neurons, because Tle4^+^ neurons are upregulated and occupied a larger area of cortical expanse in mutant mice (Fig 1E and Appendix Fig S2C). Indeed, we found that Cux1 and Ctip2 protein levels were visibly reduced in the P3 Ash2l ablation cortex (Appendix Fig S2D). At P7, we found that the neocortex was significantly thinner, with decreased upper layer marker expression, but the expression of layer VI marker Tle4 and the astrocyte marker BLBP were comparable between controls and mutant mice, indicating that there was no effect on the production of astrocytes and the layer VI cells (Appendix Fig S2A and S2B). These findings implied that the deletion of Ash2l in mice disrupted both the population and lamination of cortical PNs. Although cortical lamination as a whole was disorganized in the Ash2l cKO mice, the migration ability of upper layer PNs was dramatically compromised, whereas deep layer PNs migrated normally.

**Figure 1.**
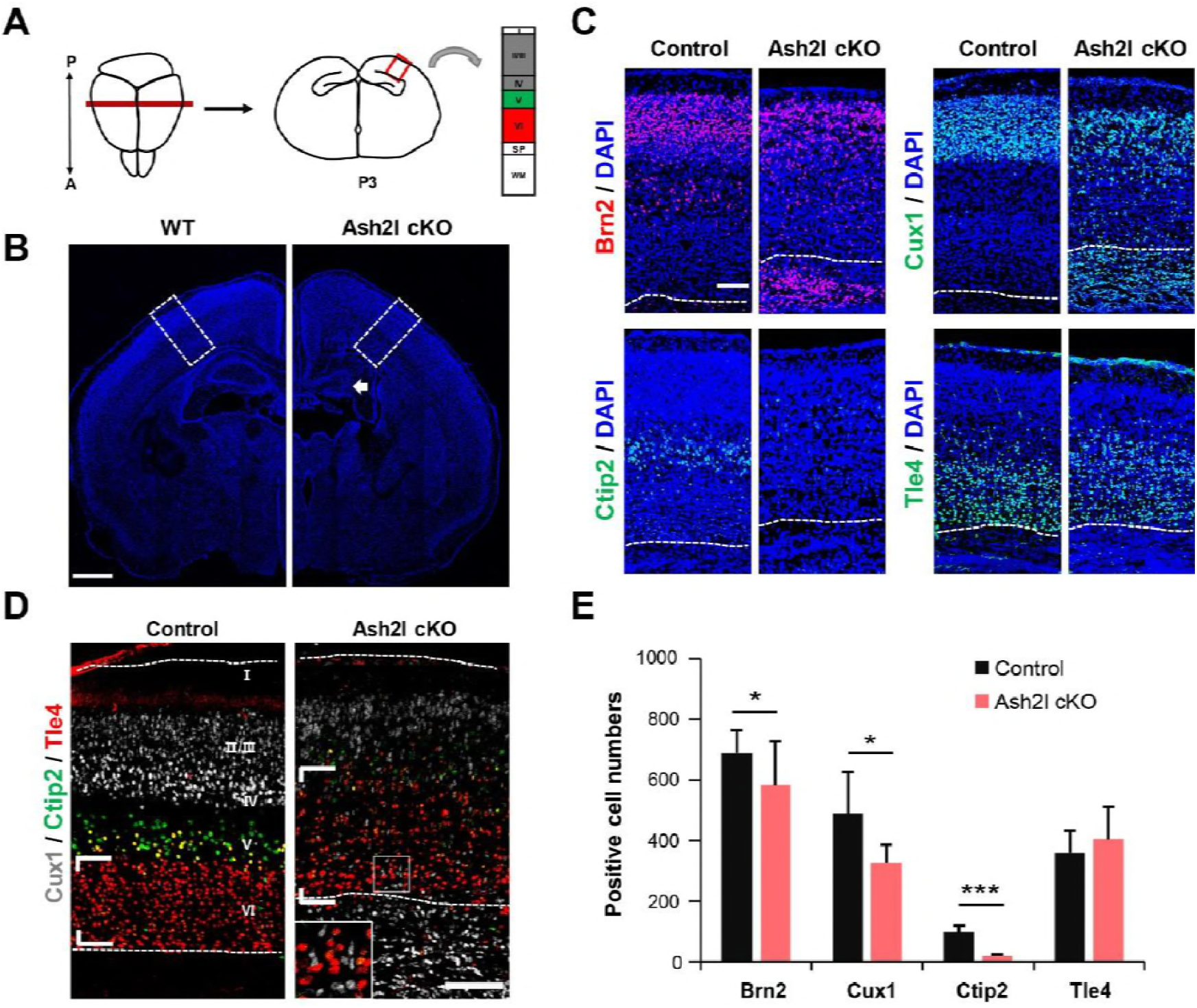
Deletion of Ash2l causes defects in the laminar architecture and cortical PN specification in the postnatal cortex. A Schematic shows area of interest (red box) in a P3 mouse coronal section, as depicted in c and e, which defines the region analysed in all postnatal experiments. A, anterior; P, posterior; SP, subplate; WM, white matter. B Immunostaining of a P3 cortex. Regions in white boxes are analysed in c and e. Arrow reveals that the hippocampus is underdeveloped in the mutant. Scale bars, 500 μm. C Immunostaining of cortical PN markers Cux1, Brn2, Ctip2 and Tle4 in the P3 neocortex. D Triple immunostaining against Cux1, Ctip2 and Tle4 in the P3 cortex. Inset shows that many Cux1^+^ cells were located in deep layers and did not co-express Ctip2 or Tle4 in the Ash2l cKO neocortex. E Quantification of Cux1^+^, Brn2^+^, Ctip2^+^, and Tle4^+^ neurons in the P3 neocortex. Regions quantified are defined as shown (B). Scale bars, 100 μm. Cux1, n=6; Brn2, n=6; Ctip2, n=8; Tle4, n=6. **p<0.01, ***p<0.001, Student’s t test. Bar graphs indicate means ± SD.

### The Late-born Upper Neurons were Decreased and Misplaced in the Ash2l cKO Neocortex

The laminar subtype of neurons is generated in a stereotyped timing schedule; most layer VI neurons are born at E12.5, followed by layer V neurons (E13.5), layer IV neurons (E14.5) and more superficial layer neurons at E15.5 and E16.5 in the mouse brain(Molyneaux, Arlotta et al., 2007). To determine whether the loss of upper layer neurons and the misspecification of PNs represented problems in the timing of PN differentiation, we conducted a series of birthdating experiments via 5-ethynyl-2-deoxyuridine (EdU) treatment, a thymidine analogue that labels proliferating cells, at E12.5, E14.5, or E16.5, followed by examination of their neuronal progenies at P3. The density of EdU-labelled neurons was comparable in control and Ash2l cKO P3 brains treated at E12.5 but significantly decreased in Ash2l cKO P3 brains treated with EdU at E14.5 and E16.5 (Fig 2A-2D). A lower proportion of EdU^+^ cells were co-stained with Ctip2 (Fig 2B and 2E), but more were co-stained with Tle4 (Appendix Fig S3A) in mice that were EdU treated at E12.5. This suggested that the Ash2l cKO mice did retain early neurogenesis ability although Ctip2^+^ neurons were reduced. EdU-positive neurons localized in deep cortical layers also maintained Cux1 immunoreactivity in the E14.5 EdU-treated Ash2l cKO cortex (Fig 2B). This indicated that the majority of EdU^+^ cells at E14.5 typically give rise to upper cortical PNs, similar with the control group.

**Figure 2.**
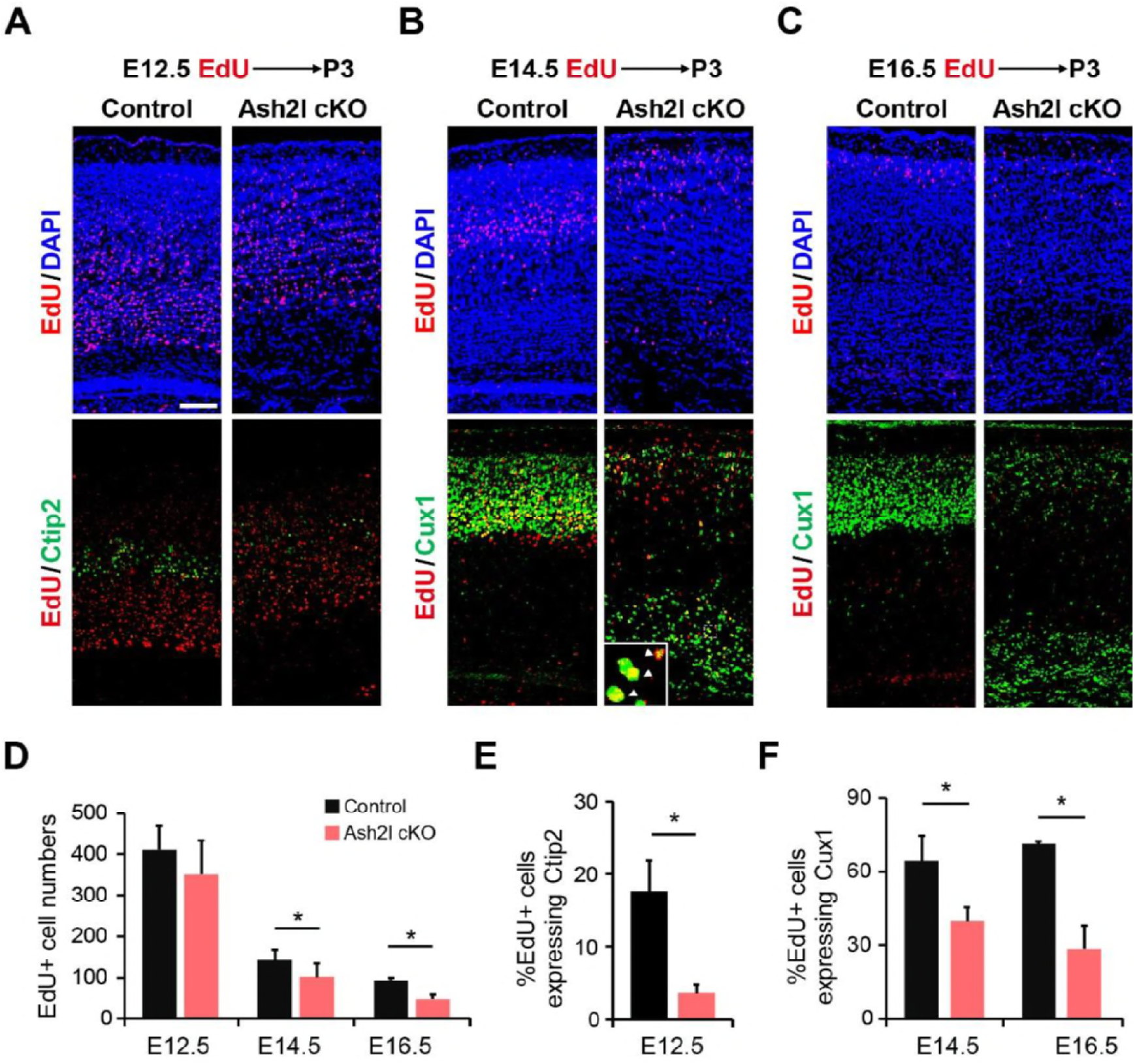
The production and migration of upper neurons were dysregulated in the Ash2l cKO neocortex. A EdU and Ctip2 immunostaining in the P3 cortex after EdU treatment at E12.5. B EdU and Cux1 immunostaining in the P3 cortex after EdU treatment at E14.5. Inset shows EdU^+^ neurons localized in deep cortical layers, which also maintained Cux1 immunoreactivity. C EdU and Cux1 immunostaining in the P3 cortex after EdU treatment at E16.5. D Quantification of EdU^+^ neurons in the P3 neocortex after EdU treatment at E12.5, E14.5 and E16.5. E Quantification of EdU^+^ and Ctip2^+^ neurons in the P3 neocortex after EdU treatment at E12.5. F Quantification of EdU^+^ and Cux1^+^ neurons in the P3 neocortex after EdU treatment at E14.5 and E16.5. Scale bars, 100 μm. n=3; *p<0.05, Student’s t test. Bar graphs indicate means ± SD.

Notably, from the EdU birthdate tracing results, the later-generated upper layer cortical neurons were located beneath the early-born deep layer neurons. This result is consistent with the previously described immunostaining data, which implied that the cortical superficial layer neurogenesis was defective and accompanied by neuronal migration or position disorders. Because of Reelin-deficient mice mutant reeler exhibited neuronal migration defects and inverted cortical layering phenotype(Frotscher, 2010, Jossin, 2004), we performed Reelin immunostaining at E12.5 and E16.5 to check whether Reelin was dysregulated after Ash2l deletion. Compared to the wild-type control cortex, the abundance and location of Reelin was unaltered in Ash2l cKO mice (Appendix Fig S3B). Therefore, it is unlikely that these neurons were incapable of migrating into the cortical plate due to abnormalities in Cajal-Retzius (CR) cell development.

Additionally, it is also interesting to note that although these neurons were capable of being generated and expressing corresponding markers, they were not in a normal state and ultimately failed to survive in the postnatal neocortex. The thickness of the cortex and the number of neurons were only slightly decreased at P3, but the cortex was significantly thinner (only half) at P7. Using TUNEL assays, there was a significant increase in TUNEL^+^ cells in the P3 Ash2l cKO neocortex (Appendix Fig S3C), which indicates the failure of cortical PNs to survive postnatally and explains the progressive reduction in cortical thickness observed at P7. Collectively, these observations indicated that deletion of Ash2l perturbed upper neuronal generation and migration, and resulted in unusually specified PNs that ultimately failed to survive in the postnatal neocortex.

### Reduction of Progenitors in the Ash2l cKO Embryonic Dorsal Telencephalon

The decreased cortical size and thickness in Ash2l mutant cortices may be due to impaired cell survival, reduced proliferation, or enhanced NPCs differentiation. Given the profound effects of Ash2l on neuronal production, we next checked the expression of NPC markers in control and Ash2l cKO cortex. As corticogenesis progresses, RGCs undergo asymmetric neurogenic divisions to self-renew and generate IPs at the ventricular surface of the VZ(Noctor, Martinez-Cerdeno et al., 2004, Stancik, Navarro-Quiroga et al., 2010). IPs localized above SVZ (the basal layer of the VZ) at later stages of neurogenesis will undergo one or more symmetric divisions before terminally differentiating to produce a pair of (or four) neurons that will largely populate the upper layers of the neocortex(Kowalczyk, Pontious et al., 2009). Compared with the control group, decreased expression of the RGC marker Pax6 was observed by immunostaining in the Ash2l cKO cortex at both E14.5 (Fig 3A and 3C) and E16.5 but it was only significantly altered at E16.5 (Fig 3D and 3F). In addition, the population of IPs identified by Tbr2 staining was significantly reduced by Ash2l depletion at both E14.5 and E16.5 (Fig 3B, 3C, 3D and 3F).

**Figure 3.**
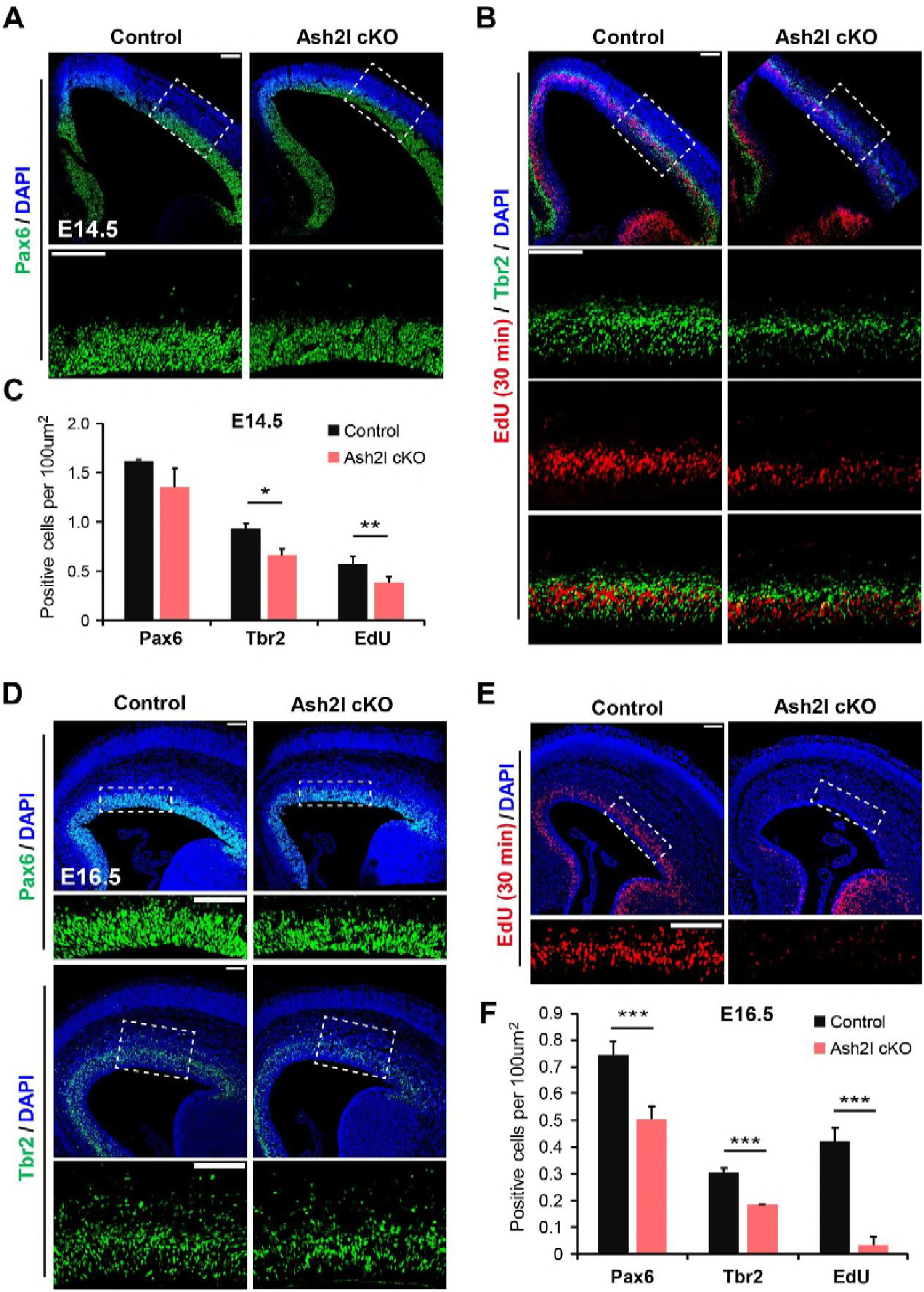
Ash2l regulates the cortical NPCs population. A Immunofluorescence staining of the RGC marker Pax6 in the E14.5 neocortex. B IP marker Tbr2 and EdU immunostaining in the E14.5 cortex. The embryos were collected 0.5 h after EdU injection. C Quantification of Pax6^+^, Tbr2^+^ and EdU^+^ progenitors in the E14.5 neocortex. D Immunofluorescence staining of the progenitor markers Pax6 and Tbr2 in the E16.5 neocortex. E EdU immunostaining in the E16.5 cortex. The embryos were collected 0.5 h after EdU injection. F Quantification of Pax6^+^, Tbr2^+^ and EdU^+^ progenitors in the E16.5 neocortex. Lower panels: higher-magnification images of areas indicated by white boxes. Scale bars, 100μm. n=3; *p<0.05, **p<0.01, ***p<0.001, Student’s t test. Bar graphs indicate means ± SD.

To further assess the proportion of proliferative cells in S phase, we injected EdU into E14.5 and E16.5 pregnant females and harvested the embryos 0.5 h after EdU pulse. The number of EdU -labelled cells were also reduced in Ash2l cKO mice (Fig 3B, 3C, 3E and 3F). Compared to E14.5, EdU^+^ cells had almost completely disappeared at E16.5 (Fig 3E). Interestingly, the numbers of Pax6 positive RGCs and Tbr2 positive IPs were unaffected at E12.5 (Appendix Fig S4A and S4B). These findings suggested that RGC and IP populations are significantly affected at late neurogenesis in Ash2l knockout mice, consistent with the above observation that upper neurons generated after E14.5 were largely disrupted in the Ash2l cKO mice.

### Abnormal Proliferation of Progenitors in the Ash2l cKO Embryonic Dorsal Telencephalon

Next, we explored the cause of the reduction in RGC/IP populations after Ash2l depletion. Immunostaining followed by quantification of the cell proliferation marker phospho-histone H3 (pH3) was performed in both Control and Ash2l deletion developing cortices. Significant reduction in basally dividing pH3 cells was observed in the SVZ regions at E14.5 (Fig 4A and 4C). There was no difference in the apical pH3-positive cell numbers in VZ regions between knockout mice and controls at E14.5 (Fig 4B), indicating that depletion of Ash2l significantly affects proliferation of IPs at the late neurogenic stage. In addition, as pH3 labels the M phase cells localized basal of the SVZ and apical of the VZ, there is a greater proportion of pH3^+^ cells was observed in the subapical VZ at E16.5 (Fig 4D and 4E). The dramatically disorganized expression pattern of apical pH3 suggested that the NPC cell cycle dynamics may altered after Ash2l deletion at the late neurogenic stages.

**Figure 4.**
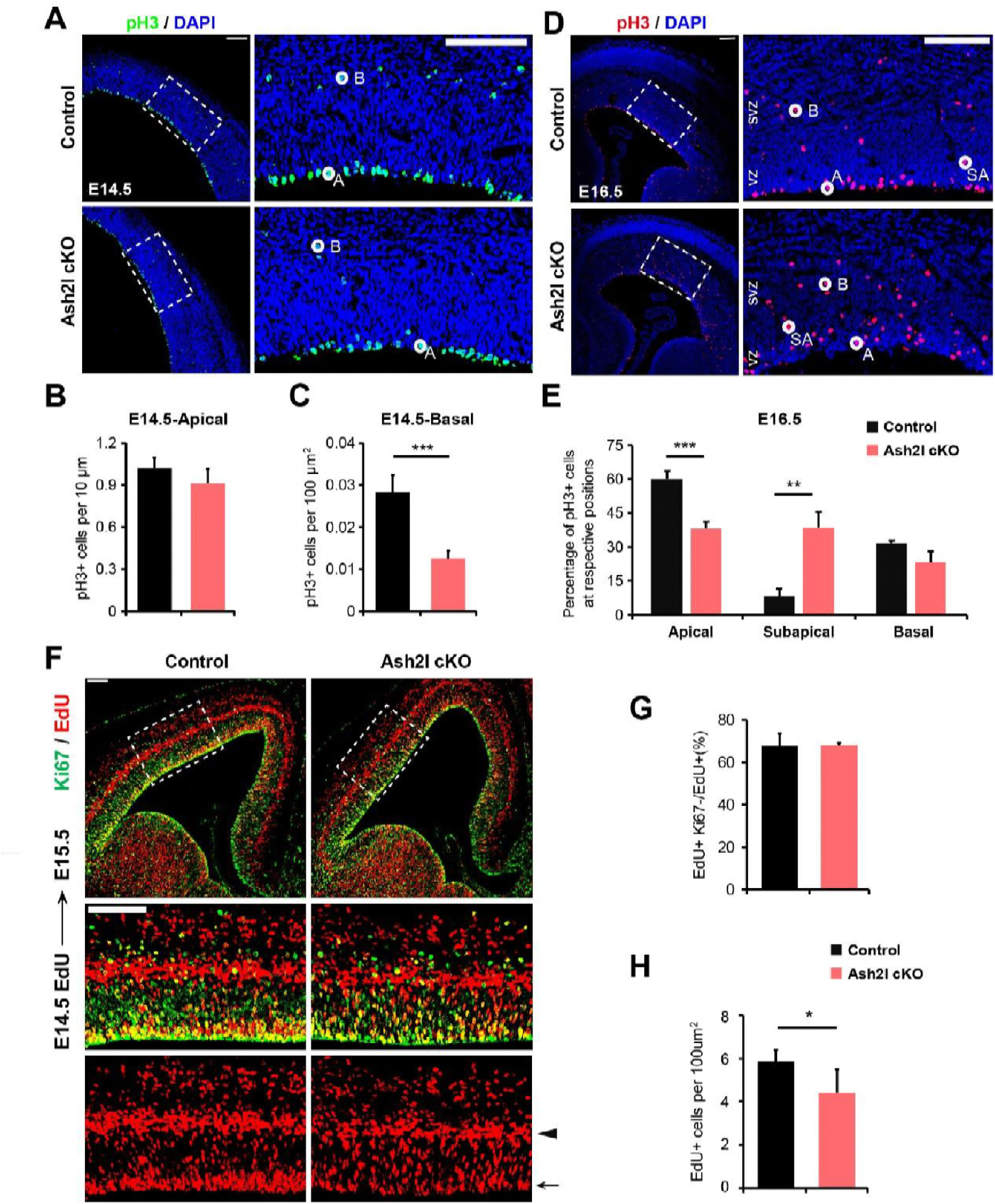
Ash2l regulates cortical NPCs proliferation during development. A Immunostaining of pH3^+^ cells in the E14.5 cortex. Left panels: higher-magnification images of areas indicated by white boxes. B Quantification of pH3^+^ cells localized in apical regions of the E14.5 cortex C Quantification of pH3^+^ cells localized in basal regions of the E14.5 cortex. C Immunostaining of pH3^+^ cells in the E16.5 cortex. Left panels: higher-magnification images of areas indicated by white boxes. D Quantification of the proportion of pH3^+^ cells localized in apical, sub-apical and basal regions of the E16.5 cortex. E Cell cycle exit analysis. Ki67 and EdU immunostaining in the E15.5 cortex after EdU treatment at E14.5. Lower panels: higher-magnification images of areas indicated by white boxes. F Quantification of the proportion of Ki67^+^ and EdU^-^ cells in the E15.5 cortex after EdU treatment at E14.5. G Quantification of the density of EdU^+^ cells in the E15.5 cortex after EdU treatment at E14.5. Arrowhead indicates SVZ, arrow indicates VZ. H SVZ, Subventricular zone; VZ, ventricular zone. A, apical; SA, sub-apical; B, basal. Scale bars, 100 μm. n=3; *p<0.05, **p < 0.01, ***p < 0.001, Student’s t test. Bar graphs indicate means ± SD.

To further confirm the differentiation ability of Ash2l cKO NPCs, we performed cell cycle exit analysis in control and Ash2l cKO brains at E15.5 and cortices were analysed 24 h after EdU incorporation, and Ki67 was used to identify cells that entered the following cell cycles. We found that the ratio of differentiated neurons was not different between Ash2l cKO and control mice but markedly reduced EdU^+^ cells, especially of VZ and SVZ, were observed in the Ash2l cKO cortex (Fig 4F-4H). This finding suggested that the proliferation and not the differentiation ability of RGCs/IPs was altered after Ash2l deletion during development. We found irregular cell death in the E16.5 Ash2l cKO cortex but not at E14.5 (Appendix Fig S4C). Furthermore, the neurosphere populations cultured *in vitro* from both control and Ash2l cKO E14.5 cortices were compared and were significantly reduced upon Ash2l disruption (Appendix Fig S4E). We also examined cell cycle alteration through flow cytometric DNA analysis; there were fewer cells in S and G2/M phase in Ash2l cKO neurospheres (Appendix Fig S4F). This finding indicated that the reduced numbers of proliferating cells after Ash2l deletion may result from disrupted cell cycle of neural progenitors. Both the *in vivo* and *in vitro* observations indicated that RGCs/IPs were unable to self-renew and proliferate normally at the late stages of corticogenesis in Ash2l knockout mice.

### Ash2l Acts as a Transcriptional Activator During Cortical Development

To gain further insight into the molecular causes of these observed phenotypes, we analysed the direct target genes of Ash2l by mapping genome-wide occupancy of ASH2L using chromatin immunoprecipitation with sequencing (ChIP-seq). Previous studies indicate that Ash2l works by mediating H3K4me3 of target genes, so we performed Ash2l and H3K4me3 ChIP-seq in cells from the neocortex at E14.5, at which point the Ash2l cKO mice began to exhibit abnormalities. Similar to previous findings from cell lines(Wan et al., 2013), our twice-repeated, *in vivo* Ash2l ChIP-seq analysis revealed that Ash2l is highly enriched in gene promoter regions and 5’UTRs (Fig 5A and Appendix Fig S5A). It is also evident from the parallel H3K4me3 ChIP-seq that Ash2l binding and H3K4me3 enrichment share an almost identical profile in gene bodies (Fig 5C). Statistical analysis showed that the vast majority (82.4%) of the Ash2l-occupied regions were also marked with significant levels of H3K4me3 and, conversely, that about half (55%) H3K4me3-enriched regions were occupied by Ash2l (Fig 5B). Strikingly, the loss of Ash2l resulted in a global reduction of H3K4me3 active marks in promoters (Fig 5D). Western blot analysis also confirmed the downregulation of H3K4me3 after Ash2l knockout (Appendix Fig S5B). Taken together, our Ash2l and H3K4me3 ChIP-seq results from WT and cKO cortices demonstrated that Ash2l could regulate tri-methylation of H3K4, a transcriptionally active mark during cortex development.

**Figure 5.**
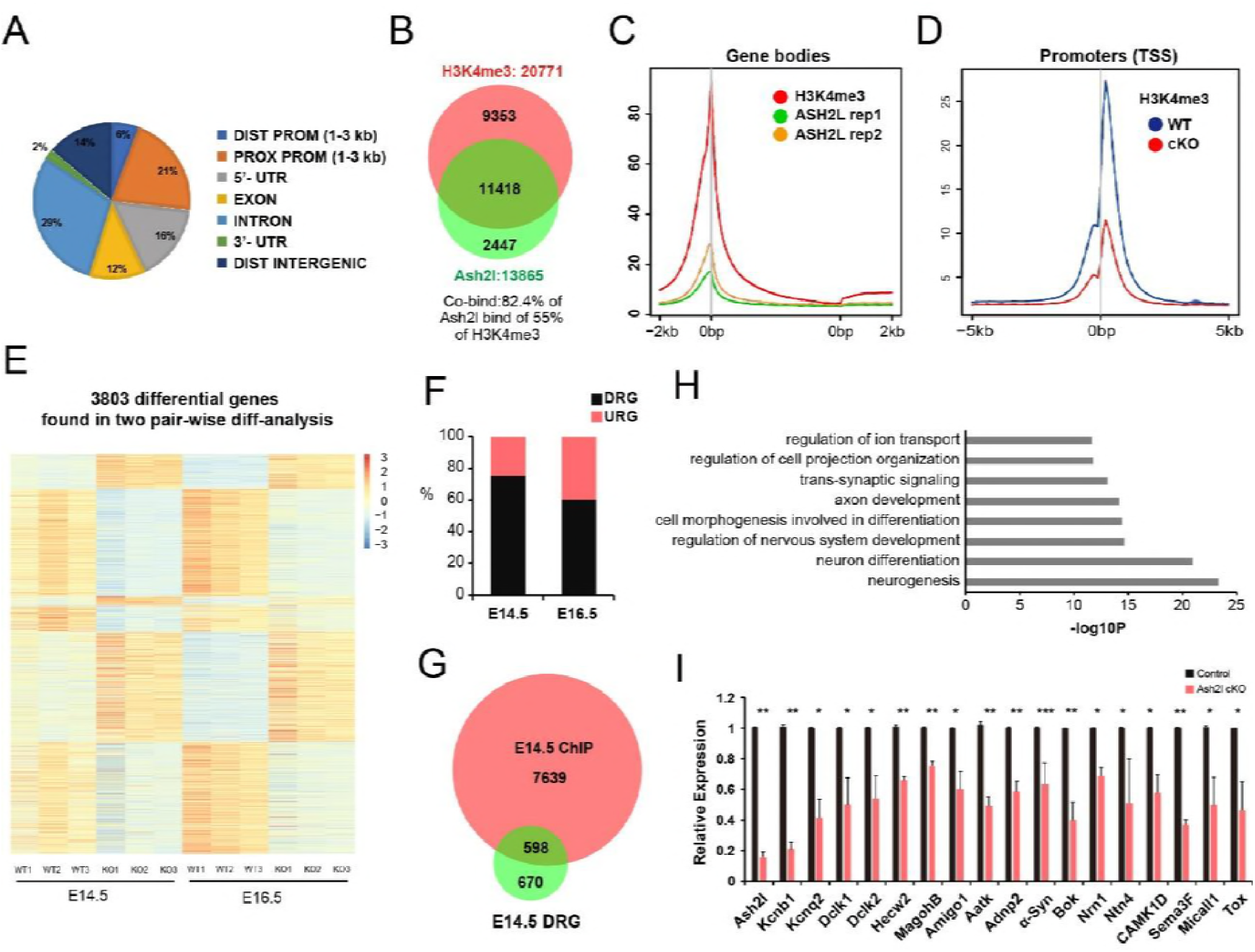
Ash2l acts as a transcriptional activator during neurogenesis. A Pie chart shows the percentage of peaks annotated to each segment in genome-wide scale in Ash2l ChIP-seq (rep1) results using WT neocortex at E14.5. B Venn diagram showing the overlap of genes occupied by Ash2l and genes enriched with H3K4me3 at high confidence in the cortex as determined by ChIP-seq. C Composite profiling of Ash2l (rep1 and rep2) and H3K4me3 ChIP signals around gene bodies as determined by ChIP-seq. D Genome-wide decreases in H3K4me3 upon deletion of Ash2l in E14.5 mutant cortices are shown, as determined by ChIP-seq analysis of mutant and control chromatin. E Gene expression cluster analysis of differentially expressed genes upon Ash2l deletion from E14.5 and E16.5 dorsal cortices from a total of 3,803 differentially expressed genes. F Histogram displays that most differentially expressed genes were downregulated at both E14.5 and E16.5. G Venn diagram showing the overlap of genes bound by Ash2l and downregulated transcripts upon Ash2l deletion at E14.5. H GO analysis of Ash2l directly regulated genes. I RT–qPCR analysis of neuronal genes in control and Ash2l cKO neocortex at E14.5 stage. n=3; *p<0.05, **p < 0.01, ***p < 0.001, Student’s t test. Bar graphs indicate means ± SD.

To further dissect the role of Ash2l in NPCs proliferation of late-neurogenesis stage, we evaluated the genome-wide analysis of gene expression patterns changes using RNA sequencing (RNA-seq) with three biological replicates from E14.5 and E16.5 cortices (Fig 5E). Ash2l deletion resulted in highly reproducible changes in the transcriptome of cortices, with 1353 genes whose expression showed a significant change in both E14.5 and E16.5 seq-data (Appendix Fig S5C). Comparing gene expression levels in WT and cKO samples at the same developmental stage, depletion of Ash2l predominantly resulted in downregulated expression of genes, which was consistent with its function as a transcriptional activator (Fig 5F). From combined analysis of Ash2l binding peaks and downregulation of transcripts after Ash2l deletion at E14.5, we identified 598 genes that were directly regulated by Ash2l during neurogenesis (Fig 5G). Gene ontology (GO) analyses of these targets revealed significant enrichment of neural development-related categories (Fig 5H). To validate these findings, we performed quantitative RT–PCR analysis of selected genes, and confirmed that numerous of neuronal genes and related elements important for neurogenesis were downregulated in the Ash2l cKO dorsal cortex (Fig 5I). We also found that a considerable number of genes involved in cell proliferation were dysregulated from E14.5 cortices RNA-seq data (Appendix Fig S5D). Together, our results show that Ash2l acts as a transcriptional activator to regulate gene expression program that necessary for cortical development.

### Ash2l Controls NPCs Proliferation by Activating Wnt-β-catenin Signalling Activity

Previous work has suggested that erased β-catenin signalling inhibits cortical NPCs proliferation and impairs radial migration of neurons at late neurogenesis(Machon, van den Bout et al., 2003). We found that genes with Ash2l binding peaks were mostly involved in the Wnt signalling pathway (Fig 6A), as well as genes (Fig 5G) that were downregulated in both E14.5 and E16.5 cortices (Appendix Fig S5E). Moreover, detailed investigation of the binding profiles revealed that Ash2l occupied the core promoter regions of Ctnnb1, Fzd7 and Sfrp2 (Fig 6B), as well as the promoters of Axin1 and Lrp1 (Appendix Fig S5F), all of which are important for the Wnt signalling pathway. To understand whether Ash2l directly targets these genes, we performed Ash2l ChIP-qPCR experiments in the E14.5 cortex and found that Ash2l can directly target the promoter regions of these genes (Appendix Fig S5G). In addition, RT-qPCR analysis also showed reduced expression of several Wnt signalling related genes (Fig 6C). To further address the hypothesis that Wnt signalling activity was downregulated after Ash2l deletion, we detected the protein level of β-catenin-the critical component of Wnt signalling pathway and found that it was downregulated, as well as Wnt downstream target gene CyclinD1 (Fig 6D). Taken together, these results suggest that Ash2l may directly regulate Wnt signalling activity during corticogenesis.

**Figure 6.**
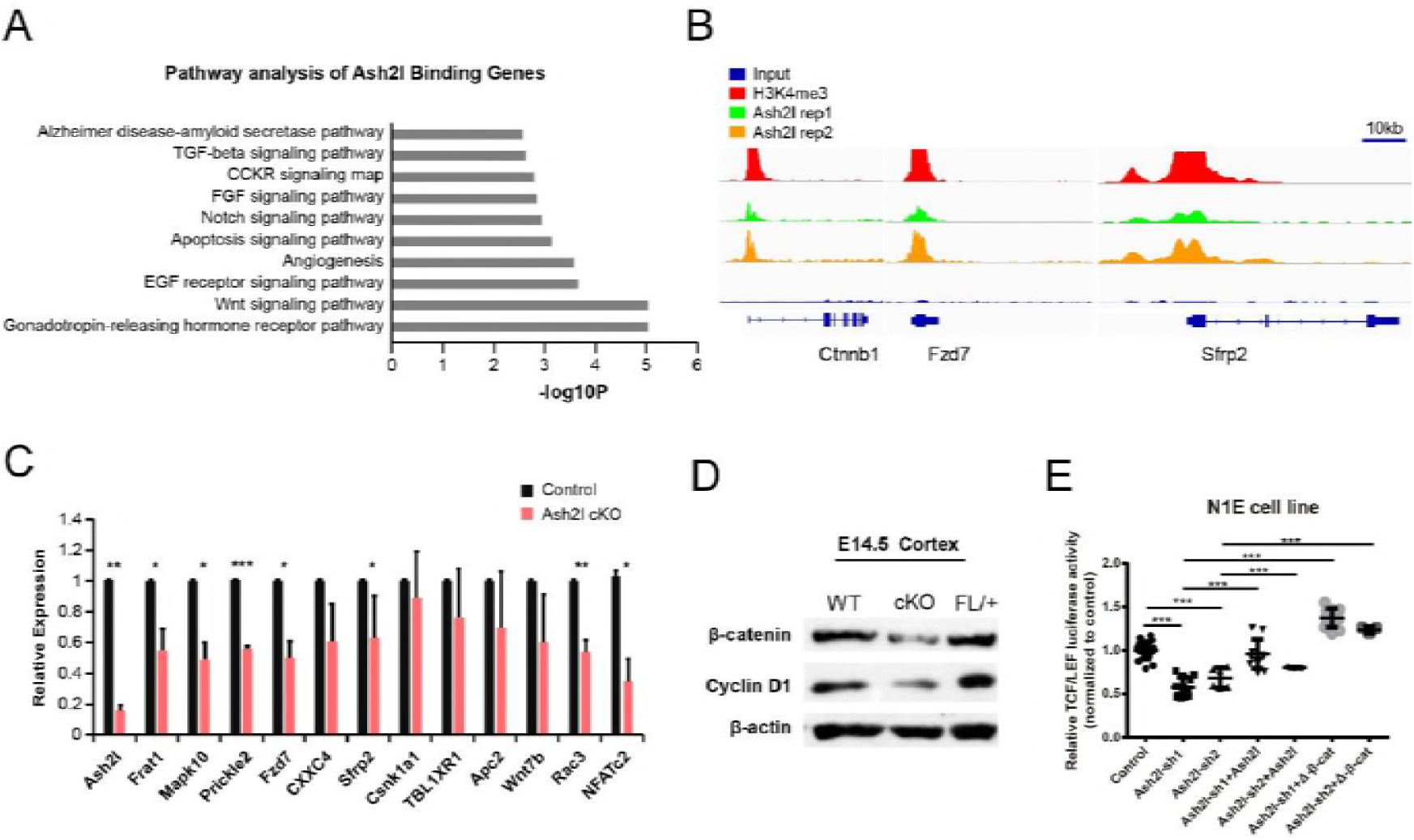
Deletion of Ash2l suppresses Wnt signalling activity. A GO analysis of Ash2l target genes reveals that one of the top enriched pathways was Wnt signalling pathway. B Gene tracks of Ash2l and H3K4me3 enrichments by ChIP-seq analysis at core promoter regions of Ctnnb1, Fzd7 and Sfrp2, which are closely related to Wnt signalling. C RT–qPCR analysis of Wnt pathway-related transcripts in control and Ash2l cKO neocortex at E14.5 (n=3; *p<0.05, **p < 0.01, ***p < 0.001, Student’s t test. Bar graphs indicate means ± SD.). D Western blot analyses of β-catenin (core factor of Wnt signalling) and CyclinD1 (target of Wnt signalling). E TCF/LEF luciferase assay in the N1E cell line using both Ash2l shRNAs and rescue with mAsh2l and ΔN-β-catenin (Control n=19, Ash2l-sh1 n=12, Ash2l-sh2 n=6, Ash2l-sh1+Ash2l n=12, Ash2l-sh2+Ash2l n=3, Ash2l-sh1+ΔN-β-catenin n=12, Ash2l-sh1+ΔN-β-catenin n=3; ***p < 0.001, one-way ANOVA followed by Bonferroni’s multiple comparison test. Bar graphs indicate means ± SEM).

To further test the above conclusion, we performed *in vivo* reporter assays through electroporating the TOP-dGFP construct containing a destabilized GFP under the control of β-catenin/TCF binding sites. We electroporated TOP-dGFP reporter plasmids plus a Cre-mCherry-expressing plasmid (pCI-Cre) into the E14.5 Ash2l^fl/fl^ embryos and tissues were collected at E16.5 and processed for immunofluorescence analysis (Fig 7A). Compared with the wild controls, eliminating Ash2l significantly reduced the ratio of GFP^+^ cells, both progenitors and neurons, and this effect was rescued by co-expressing mAsh2l alongside the pCI-Cre construct (Fig 7B and 7C). Similarly, silencing of Ash2l markedly reduced Wnt signalling activity in N1E-115 cells *in vitro*, and both mAsh2l and ΔN-β-catenin (a degradation-resistant β-catenin mutant construct) could restore this effect (Fig 6E and Appendix Fig S5H). Taken together, our *in vivo* and *in vitro* observations prove that Ash2l, as a transcriptional activator, regulated the Wnt-β-catenin signalling pathway and suggest that the primary cause of the defective proliferation of Ash2l cKO progenitors may depend on the Wnt-β-catenin signalling pathway.

**Figure 7.**
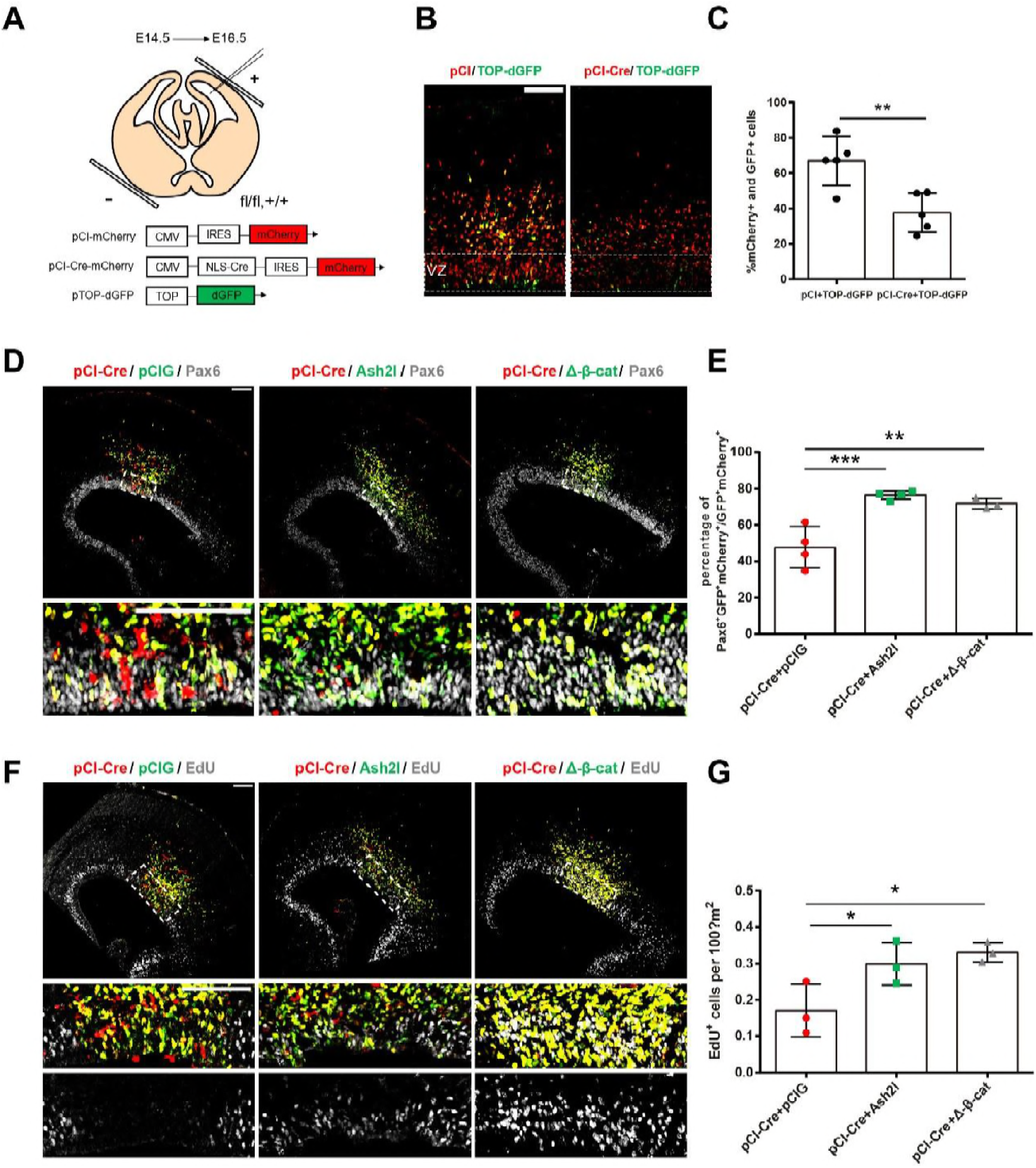
Ash2l regulates NPCs proliferation in a β-catenin-dependent manner. A Schematic of electroporation protocol and plasmids used in (B). B-C Immunofluorescence and quantitative TOP analyses in E16.5 cortices electroporated with the pCI-mCherry or pCI-Cre-mCherry construct together with TOP-dGFP at E14.5. The white box indicates VZ. VZ, ventricular zone. D-E Immunofluorescence and quantitative analyses of Pax6 in E16.5 cortices electroporated with pCI-Cre plus pCIG, mAsh2l and ΔN-β-catenin plasmid. The Pax6^+^ cells of VZ were quantified. Embryos were collected half an hour after EdU injection. (n=3) F-G Immunofluorescence and quantitative analyses of EdU in E16.5 cortices electroporated with pCI-Cre plus pCIG, or mAsh2l or ΔN-β-catenin plasmid. The EdU^+^ cells of VZ were quantified. Embryos were collected half an hour after EdU injection. (pCIG, n=4; mAsh2l, n=4; ΔN-β-catenin, n=3) Lower panels: higher-magnification images of areas indicated by white boxes. Scale bars, 100 μm. *p<0.05, **p < 0.01, ***p < 0.001, one-way ANOVA followed by Bonferroni’s multiple comparison test. Graphs indicate means ± SEM.

To functionally test whether the increased Wnt signalling activity could also rescue progenitor proliferation defects caused by Ash2l knockout, we next performed *in utero* electroporation in Ash2l^fl/fl^ embryos with pCI-Cre vector, and co-induced with ΔN-β-catenin plasmid that could consistently activate Wnt-β-catenin activity. First, we found that, similar to the E16.5 Ash2l cKO mice cortex, 30-min EdU labelling cells number was visibly reduced in the VZ of E16.5 Ash2l^fl/fl^ cortices that underwent *in utero* electroporation with pCI-Cre vector at E14.5 compared to those that underwent electroporation with the pCI vector (Appendix Fig S6). Next, we directly determined whether Ash2l regulates cortical development via activation of Wnt-β-catenin signalling. We electroporated ΔN-β-catenin together with pCI-Cre vector in E14.5 Ash2l^fl/fl^ cortices and found that the number Pax6^+^ cells increased in E16.5 cortices, similar with the electroporation results of mAsh2l with the pCI-Cre vector did (Fig 7D and 7E). Furthermore, ΔN-β-catenin also rescued progenitors proliferation ability, we observed a significant increase of EdU^+^ (0.5 h) cells in the VZ when electroporated together with pCI-Cre vector in E14.5 Ash2l^fl/fl^ cortices (Fig 7F and 7G). It means that β-catenin can rescue the progenitor proliferation defects caused by Ash2l deletion. Hence, our results demonstrate that Ash2l regulates late neurogenesis in a Wnt-β-catenin dependent manner. These findings suggest that COMPASS family epigenetic factor Ash2l is required for proper cortical development through appropriate activation of Wnt signalling at the late neurodevelopmental stages.

## Discussion

### Ash2l-Dependent Histone Modification is Essential for Maintaining the Cortical NPC Statement

Neurogenesis is a highly organized process with timely specification of the NPCs lineage; it relies on the specific transcriptional program to control extremely precise gene expression. Generally, during the cell-type-specific fate acquisition and differentiation, the related genes are marked with histone H3K4me3 when actively transcribed and marked with H3K27me3 in the repressed state or switch their “bivalent” modification signature through the modification of their promoters. COMPASS and PcG proteins are highly evolutionarily conserved lysine methyltransferases that regulate the methylation of H3K4 and H3K27, thereby activating or silencing gene expression, respectively(Piunti & Shilatifard, 2016, Schuettengruber et al., 2017). However, the role of COMPASS proteins remains largely unknown. In this study, we chose ASH2L, one of the four well-conserved core subunits essential for modifying activities, to reveal that COMPASS complexes are essential for the maintenance of a proper differentiation program of NPCs during development.

We generated mice lacking Ash2l specifically in neural stem cell (NSC) of the dorsal telencephalon at early stages of cortical development, as the NPC that produces all neocortical projection neurons. Our findings reveal that genetically removing Ash2l resulted in NPCs exhaustion and generation of abnormal cell progeny during late cortical neurogenesis. After conditionally knock out Ash2l, the following changes occurred in NPCs: (1) reduced NPC numbers begin at E14.5 (Fig 3); (2) decreased RGC proliferation, especially at E16.5 (Fig 3E); (3) increased progenitor cell death (Appendix Fig S4C); (4) change in RGC cell cycle kinetics (Appendix Fig S4F); and (5) reduction of progeny IPs and PNs (Fig 3 and 2). Thus, Ash2l is necessary to maintain the proper number and state of NPCs, which is essential to ensure correct brain development. Notably, COMPASS complex and H3K4 tri-methyltransferase have been reported that participate in the coordination of cell cycle progression(Beilharz, Harrison et al., 2017, Tajima, Yae et al., 2015). Jiao and colleagues found that HIRA can recruit the H3K4 tri-methyltransferase Setd1A to maintain NPCs proliferation and terminal mitosis(Li & Jiao, 2017). These reports are highly consistent with our findings.

Aberrations in NPCs proliferation and differentiation are reasons for many neurodevelopmental disorders. In fact, malformations of the cerebral cortex are a cause of mental retardation and epilepsy, resulting from defects in neuronal migration or NPCs proliferation(Barkovich & Kuzniecky, 2000, Olson & Walsh, 2002). Therefore, the root cause of a range of phenotypes, including retarded growth, epileptic behaviour, neonatal death, hydrocephalus, and cortical lamination defects, in the Ash2l conditional ablation mouse may be the destruction of the NPCs fate decision program. Moreover, some ion channel proteins are abnormally expressed (Fig 5H and 5I), which may due to including abnormal neuronal maturation as well as abnormal expression of those proteins in the NPCs.

### Ash2l Maintains the State of NPCs in a Stage-Dependent manner

Interestingly, Ash2l maintains the proliferation and differentiation of NPCs in a stage-dependent manner. Compared with control mice, the Ash2l cKO brain showed more significantly serious defects at the late stages of neurogenesis than at the early stages, as follows: (1) Pax6^+^ RGCs were significantly reduced at E16.5 but slightly changed at E14.5(Fig 3A and 3D); (2) the single-pulse, short-time EdU incorporation experiment showed that the reduction of EdU^+^ cells at E16.5 was much more severe than at E14.5 (Fig 3B and 3E); (3) the EdU birthdating assay showed that the late-born Brn2^+^ or Cux1^+^ upper cortical PNs were reduced and did not arrive at the correct position of the cortex, but the early-born neurons had no obviously changes (Fig 2); (4) cell apoptosis became evident only in E16.5 embryos and many downregulated genes related to “Negative Regulation of Neuron Apoptotic Process” only appeared in E16.5 cortices RNA-seq data (Appendix Fig S4C and S4D); and (5) RNA-seq data revealed much more altered expression transcripts in the E16.5 than in the E14.5 cortex after Ash2l deletion (Appendix Fig S5C).

Therefore, Ash2l deletion had markedly more impact on upper layers, which may be attributed to distinct regulatory mechanisms in RGCs and IPs such as regulating the sequential activation of stage-specific gene expression programs. This is consistent with the single cell sequence data, in which Ash2l was highly expressed in the IPs of SVZ and upper layer neurons of the cortical plate (Appendix Fig S1B); the upper layer neurons are typically derived from Tbr2^+^ IPs. Furthermore, although cortical RGCs in the Ash2l cKO neocortex continued to obtain the identity of IPs and dorsal telencephalic neurons, the RGCs could have a systematic change of cell characteristics and competency. For example, the proportion of RGCs labelled with EdU in 30 min was drastically reduced compared to their changes in number (Fig 3). Thus, RGCs were unable to preserve their stable state and the capacity of producing IPs and new-born neurons without Ash2l, which may be because of its action through the Wnt signal pathway, as discussed below.

### Ash2l Positively Regulates Wnt Signalling in NPCs

By using a combination of genome-wide DNA binding site identification (ChIP-seq), transcriptomics (RNA-seq) and functional analysis, we demonstrated that Ash2l directly binds to many targets of the Wnt signalling pathway (both canonical and noncanonical Wnt pathways), indicating that Wnt signalling might play a critical role as an effector of the observed forebrain defects after Ash2l deletion. The functions of Wnt signalling in cortical development have been extensively studied in proliferating progenitor/stem cells in the ventricular germinal zone(Chenn, 2008). It is generally believed that Wnt signalling, especially canonical Wnt-β-catenin signalling, plays a diverse role in the development of the cortex. Conditional knockout of β-catenin from NPCs by Emx1-Cre leads to neocortical malformations, reduced cell proliferation, decreased RGC proliferation, and subsequent depletion of RGC and IP progenitor pools, leading to reduced neuron numbers(Draganova et al., 2015). When used D6-Cre to mediate β-catenin conditional knockout, it results in decreased RGC proliferation after E15.5, disrupted RGC interkinetic nuclear migration, and disassembly of the radial glial scaffold impaired radial migration of neurons towards superficial layers was observed(Backman et al., 2005, Machon et al., 2003). Conversely, induction of Wnt signalling via overexpression of stabilized β-catenin increases NPCs proliferation by negatively regulating cell cycle exit and differentiation(Chenn & Walsh, 2002). Pleasure and colleagues used LRP6 mutant mice to inhibit the canonical Wnt signalling pathway and found that LRP6 mutants had a smaller, thinner cortex, as proliferation of NPCs became progressively defective at the late stages of neurogenesis(Zhou, Borello et al., 2006). Intriguingly, Machon and colleagues recently reported deletion of the Wnt co-receptor Tcf4, that efficiently blocked β-catenin signalling without disrupting adhesion, also results in reduced of RGCs proliferation and the IPs number, and hypoplastic forebrain(Chodelkova, Masek et al., 2018).

In this study, the phenotype of Ash2l cKO was very similar to that of inactive β-catenin, including a reduction in the number and proliferation of RGCs, and abnormal neuronal migration during the later stages of cortical development. In addition, the RNA-seq and RT-qPCR data of Ash2l cKO showed a number of downregulated Wnt signalling genes, and ChIP-seq and ChIP–qPCR analysis revealed that Ash2l directly occupied the core promoter regions of these genes. Furthermore, we validated that Wnt-β-catenin activity was downregulated *in vivo* through electroporating TOPFLASH reporter assays. Moreover, we used *in vivo* experiments to confirm that overexpression of β-catenin can rescue the RGC proliferation defects in the Ash2l cKO cortex. Jiao and his colleagues also identified that H3K4me3 was bound upstream of the β-catenin promoter and modulated β-catenin expression *in vitro*(Li & Jiao, 2017).

On other hand, defects in Wnt signalling have recently been strongly implicated in the development of hydrocephalus(Meng, Li et al., 2015, Ohata, Nakatani et al., 2014, Tissir, Qu et al., 2010, Xu, Xu et al., 2015), similar to the phenotype of Ash2l cKO. In addition, recent study showed that hyperpolarization of RGCs with Kir2.1 droved neuronal diversity via repression of canonical Wnt signalling(Vitali, Fievre et al., 2018), and activity-dependent modulation of canonical Wnt signalling has previously been reported to occur at the late stages of differentiation(Ataman, Ashley et al., 2008, Wayman, Impey et al., 2006). We found a series of specific potassium ion channel genes were directly targeted by Ash2l through our ChIP-seq data and their expression were downregulated, as well as Wnt signalling activity (Data not shown). We speculate that Ash2l could specially regulate the expression of potassium ion channel during neurogenesis. In addition, we cannot rule out roles for the non-canonical Wnt signalling pathway in these processes, as some noncanonical Wnt signalling transcripts were also significantly altered after Ash2l knock out (Fig 6C).

Mutation of the COMPASS member gene has been described as a cause for severe microcephaly in clinic(Yang, Baltus et al., 2012). Mutation of Ash2l are also linked to intellectual disability of infants(Karaca et al., 2015). Strikingly, disease ontology analysis of Ash2l targeted genes showed enrichment related to schizophrenia, autistic disorder and epileptic encephalopathy (Appendix Fig S5I). These fingdings deepen our understanding of the Ash2l related neurological diseases and it may be helpful for the therapeutic purposes.

Altogether, our results demonstrate that the histone modifying factor Ash2l plays a crucial role in late stages development of the mammalian cortex through selectively inducing Wnt-related gene expression in NPCs, establishing a timely and specific program to ensure the generation of appropriate numbers of NPCs and neurons at late cortical development. These also provide insights into how H3K4me3 contribute to cellular transition during neocortical development.

## Material and Methods

### Mice

Adult Ash2l-fl-neo mice were obtained from Beijing Biocytogen. Ash2l^fl/fl^ mice were generated by crossing Rosa26 Flp mice. Ash2l-cKO mice were produced by crossing Ash2l^fl/fl^ with D6-Cre mice. All mice had free access to food and water. All animal experiments were conducted according to protocols approved by the Institutional Animal Care and Use Committee at the Academy of Medical Sciences and Peking Union Medical College (PUMC).

### TUNEL assay, EdU staining, and immunofluorescence staining of brain sections

TUNEL staining was conducted on cryosections using an *in situ* Cell Death Detection Kit (11684817910, Roche). Pregnant mice at defined pregnancy stages were injected intraperitoneally with 50 ug/g body weight of EdU. EdU staining was conducted on cryosections with a Click-iT plus EdU Alexa Fluor 594 Imaging Kit (Life Technologies) according to the manufacturer’s instructions. Immunofluorescence analyses of the cryosections and cells were conducted as previously described(Zhong, Feder et al., 1996). Images were acquired using a confocal laser scanning microscope (FV1000MPE-BX61WI, Olympus) and were analysed using FluoView (Olympus) and Adobe Illustrator (Adobe Systems). Primary antibodies used were as follows: Pax6 (1:500, PRB-278P, Convance), Tbr2 (1:500, ab23345, Abcam), Brn2 (1:500, sc-6029, Santa Cruz), Cux1 (1:500, sc-13024, Santa Cruz), Tle4 (1:500, sc-365406, Santa Cruz), Ctip2 (1:500, ab18465, Abcam), Reelin (1:500, ab78540, Abcam), Ki67 (1:1000, ab16667, Abcam), and BLBP (1:1000, ab32423, Abcam). Secondary antibodies used were as follows: goat anti-mouse, anti-rabbit or anti-rat Alexa-546-, Alexa-488-and Alexa-647-conjugated antibodies (1:500, Invitrogen). DAPI (ZLI-9557, ZSBio) was used for DNA staining to reveal the nuclei.

### Image acquisition and analysis

4×4 tiled images were obtained to cover each coronal section with a 10x or 40x objective by scanning microscope and compared with equivalent sections in littermate counterparts. For quantification of labelled neurons at postnatal stages, a region of the lateral neocortex, including layers I–VI, subplate and white matter (defined in Fig 1A) was measured to determine the area. For quantification of labelled progenitors at embryonic stages, fluorescently labelled cells (Pax6^+^, Tbr2^+^, and EdU^+^ cells) adjacent to the lateral ventricles were ere quantified to measure the number of cells per 100 μm^2^. Apical pH3^+^ progenitors (Fig 4B) were quantified by measuring the number of labelled cells along the length of the ventricular surface. For quantitative analysis of electroporated neocortices, only mCherry^+^ cells localized within the dorso-lateral cortex were examined.

### Western blotting

For Western blotting analysis, cortical cells were lysed in lysis buffer (50 mM Tris-HCl, 150 mM NaCl, 0.5% Triton X-100, 1 mM EDTA, 1 mM Na_3_VO_4_, 25 mM NaF, 10 Mm Na_4_P_2_O_7_•10H_2_O) and protease inhibitor (5 μg/ml PMSF, 0.5 μg/ml leupeptin, 0.7 μg/ml pepstatin, and 0.5 μg/ml aprotinin). Proteins were separated by SDS–PAGE and transferred onto a nitrocellulose membrane. After membranes were blocked with 5% milk for 30 min, they were probed with various primary antibodies overnight at 4°C, followed by incubation with secondary antibodies for 2 h at room temperature and visualization with enhanced chemiluminescence reagent (Thermo Scientific). Antibodies included Ash2l (1:1000, A300-489A, Bethyl Laboratories), H3k4me3 (1:1000, 07-473, Abcam), H3k27me3 (1:1000, 9733, Cell Signaling Technology), H3K36me3 (1:1000, ABE305, Millipore), H3 (1:1000, 06-755, Millipore), Cux1 (1:500, sc-13024, Santa Cruz), Ctip2 (1:800, ab18465, Abcam), CyclinD1 (1:500, sc-8396, Santa Cruz), β-catenin (1:1000, 8814, Cell Signaling Technology), and β-actin (1:5000, A5441, Sigma).

### Reverse transcriptase–quantitative PCR

Total RNAs of control and Ash2l cKO brains at desired developmental stages were extracted using TRIzol reagent (Invitrogen) according to the manufacturer’s instructions. Complementary DNA synthesis was performed with the Reverse Transcriptase Kit (TransGene) following the manufacturer’s instructions. Quantitative RT–PCR was performed using a SYBR green-containing PCR kit (TaKaRa), and signals were detected with the IQ5 sequence detection system (Applied Biosystems). Related primer sequences are listed in the **Appendix Table 1**, and Gapdh was used as endogenous control.

### DNA constructs and transfection

Ash2l shRNA sequences were cloned into the pLL3.7 vector, and Ash2l sequences were cloned into pCIG vector. Super8×TOPFLASH (Addgene) and a renilla-Luc-TK reporter (pRL-TK, Promega) were used for testing TCF/LEF transcriptional activity. ΔN-β-catenin was constructed by cloning a β-catenin DNA sequence containing an NH2-terminally truncated mutation in pCIG. All plasmids were purified using the Plasmid Mini Kit (QIAGEN), and then N1E-115 cells were transfected using LipofectamineTM 2000 (Invitrogen). Ash2l-shRNA sequences were described elsewhere(Wan et al., 2013).

### Luciferase assay

N1E cells at 1 × 10^5^ cells per well density were plated into 24-well plates without antibiotics. Cells were transfected with 0.8 μg of shRNA plasmid along with 100 ng of Super8×TOPFLASH and 20 ng of pRL-TK. The media were replaced with one containing antibiotics 6 h after transfection. At 48 h after transfection, cells were collected for luciferase assay. For the rescue experiments with mAsh2l, 0.2 μg of mAsh2l was co-transfected with 0.6 μg of Ash2l shRNA. For the rescue experiments with ΔN-β-catenin, 0.1 μg of ΔN-β-catenin was co-transfected with 0.7 μg of Ash2l shRNA.

### NPC culture and cell cycle analysis

NPCs were isolated from E14.5 mice by dissection of the lateral walls of the lateral ventricles and digested into single cell suspension with Accutase (Sigma). Cells were cultured in DMEM/F12 proliferation medium supplemented with 2% B27 supplement, 20 ng/ml EGF, 20 ng/ml bFGF and 0.2% BSA for proliferation. Cells were plated at 1×10^4^/ml density in 96-well plates for quantification. The primary NPCs were harvested and assessed for cell cycle analysis using PI staining through flow cytometry (BD Biosciences) as described elsewhere(Shen, Vignali et al., 2017).

### *In utero* electroporation

The Institutional Animal Care and Use Committee of Medical Sciences Academy and Peking Union Medical College approved all experiments. Pentobarbital sodium (0.7 mg/g) was used to anaesthetize pregnant mice. Five pulses of 30 V (50 ms on/950 ms interval) were delivered across the head of the E14.5 embryos. For TOPFLASH analysis, TOP-dGFP, pCI-Cre and control were electroporated at 1.5 μg/μl. For rescue experiments with mAsh2l and stabilized ΔN-β-catenin, these plasmids were also electroporated at 1.5 μg/μl.

### RNA-seq and data analysis

For RNA-seq analysis, RNA was isolated from E14.5 and E16.5 dorsal cortices with TRIzol reagent (Invitrogen) according to the manufacturer’s instructions. RNA-seq was performed by Novel Bio Ltd, China. RNA-seq data analysis was performed by Active Motif. Read mapping statistics (e.g., number of inputs, uniquely mapped reads) were obtained from STAR (v2.5.2b)(Dobin, Davis et al., 2013). Fragment assignment statistics obtained from featureCounts in the Subread software packages(Liao, Smyth et al., 2014). Differential gene expression analysis was performed through DESeq2 (v1.14.1)(Love, Huber et al., 2014).

### ChIP-seq and ChIP–qPCR

Embryonic dorsal cortices were isolated from WT and Ash2l cKO embryos at E14.5. The H3K4me3 IP experiment using WT and cKO samples were conducted in our lab; Chromatin immunoprecipitation was performed using MAGnify Chromatin Immunoprecipitation System (Invitrogen) as elsewhere described(Blecher-Gonen, Barnett-Itzhaki et al., 2013). Briefly, 200 mg brain tissue from E14.5 mice cortices were minced and cross-linked with formaldehyde (1% final concentration) for 10 min at room temperature. Stop the cross-linking reaction by adding glycine to a final concentration of 0.15 M and washed twice with ice cold PBS. Cells were resuspended in 1× lysis buffer (lysis buffer 1, lysis buffer 2 and lysis buffer 3) supplemented with protease inhibitors, followed by sonication using a bioruptor (medium power, 30 s on/30 s off for 80 cycles). Coupling antibodies to the magnetic protein G beads for 2 h at 4 °C in parallel with cell lysis and sonication. Aliquot sonicated material into microcentrifuge tubes containing protein G beads coupled to H3K4me3 (07-473, Abcam) and IgG and tumble the tubes overnight at 4 °C. Cross-linked protein–DNA complexes were eluted from the beads using 1× direct elution buffer (add 1μl 10mg/ml RNase A) and incubated 37 °C for 1 h, followed by reversion of crosslinking with followed by reversion of crosslinking with Proteinase K (65 °C for 2 h). DNA was purified using MinElute Reaction cleanup kit (QIAGEN). Sequencing libraries were constructed with 100 ng of DNA (ChIP or input) using the NEBNext Ultra DNA library prep kit for Illumina (New England Biolabs, NEB). The Ash2l and H3K4me3 IP experiments of WT E14.5 cortices, ChIP-seq and following data analysis were performed by Active Motif Inc. Ash2l antibody (39099, Active Motif) and H3K4me3 antibody (39915, Active Motif) were used in the ChIP assays. The 75-nt sequence reads generated by Illumina sequencing were mapped to the genome using the Burrows-Wheeler Aligner algorithm with default settings. Only reads that passed Illumina’s purity filter, aligned with no more than 2 mismatches, and mapped uniquely to the mouse genome (mm10) were used in the subsequent analysis. For identifying intervals over input control, the MACS cutoff for the false discovery rate after controlling for significance was P=1×10 ^–7^. The data visualization was generated using Integrative Genomics Viewer. The E14.5 WT cortices chromatin for ChIP-qPCR assay was the same as that in the ChIP-Seq analysis prepared by Active Motif Inc. Related primer sequences are listed in the **Appendix Table 2.**

### Functional analysis and disease ontology

Functional GO enrichment analysis (Fig 5H) and Pathway enrichment analysis (Appendix Fig S5E) of genes directly regulated by Ash2l were performed using ConsensusPathDB(Herwig, Hardt et al., 2016), which calculates enrichment p values using the Wilcoxon’s matched-pairs signed-rank test. Pathway analysis of Ash2l binding genes was performed using PANTHER Overrepresentation Test(Mi & Thomas, 2009) (Fig 6A), which calculates enrichment p values using the Fisher test. Disease association analysis was performed using WebGestalt(Zhang, Kirov et al., 2005) (Appendix Fig S5I).

### *In situ* hybridization

The procedures for ISH and Ash2l probe sequence were the same as previously described(Gray, Fu et al., 2004, Zhou, Wang et al., 2017).

### Statistical analysis

All plots were generated and statistical analyses were performed using SPSS 19.0 and Graphpad Prism 6.0 software. Paired Student’s t-test were used for comparison of two data sets by using SPSS 19.0, and results are presented as mean ± SD. All quantifications were performed with at least 3 brain sections from at least 3 animals. One-way ANOVA followed by Bonferroni multiple comparison tests was used for experiments with three or more data sets (luciferase assays and rescue immunofluorescence staining analyses) and results are presented as mean ± SEM. Molecular and biochemical analyses were performed using a minimum of three biological replicates per condition. For flow cytometry analysis of cell-cycle, WT neurosphere were derived from 3 embryonic cortices and cKO neurosphere were derived from 1 embryonic cortex. Venn diagrams were performed by BioVenn.

## Expanded View Materials

Supplementary materials contain 6 appendix figures, 2 appendix tables and 1 video.

## Acknowledgements

This work was supported by grants from the National Key Research and Development Program of China (2016YFA0100702, 2016YFC0902502), National Natural Science Foundation of China (31670789, 31671316), and CAMS Innovation Fund for Medical Sciences (CIFMS, 2016-I2M-2-001, 2016-I2M-1-001, 2016-I2M-1-004, 2017-I2M-2-004, 2017-I2M-1-004).

## Author Contributions

X.P., P.S., and B.Q. initiated and designed the experiments; L.L. performed most characterization of cKO mice phenotypes and molecular experiments; X.R., P.C. and P.S. conducted *in utero* electroporation experiments. C.W., X.Z., and P.X. characterized Ash2l cKO mice line. C.W., L.Z. and Q.W. cloned mouse Ash2l construct, *ash2l* probe and Ash2l-shRNA. B.Y., L.H. and W.L. prepared cultured N1E cell line and help with isolate NPCs from mice cortices. L.L. and P.S. analysed all related data. J.Y. helped design the project. L.L., P.S. and X.P. wrote the manuscript with critical thoughts from all of the authors. All authors approved the final version of the manuscript.

